# Repeated range expansion and niche shift in a volcanic hotspot archipelago: radiation of Hawaiian Euphorbia (Euphorbiaceae)

**DOI:** 10.1101/056580

**Authors:** Ya Yang, Clifford W. Morden, Margaret J. Sporck-Koehler, Lawren Sack, Paul E. Berry

**Affiliations:** Department of Ecology and Evolutionary Biology, University of Michigan, Ann Arbor. 830 North University Avenue, Ann Arbor, MI 48109-1048, USA; Department of Botany, University of Hawaii at Manoa, 3190 Maile Way, Honolulu, HI 96822, USA; Department of Land and Natural Resources, Division of Forestry & Wildlife, 1151 Punchbowl Street Rm. 325 Honolulu, Hawaii 96813, USA; Department of Ecology and Evolutionary Biology, University of California, Los Angeles, 621 Charles E. Young Drive South, Los Angeles, CA 90095-1606, USA; University of Michigan Herbarium, Department of Ecology and Evolutionary Biology, 3600 Varsity Drive, Ann Arbor, Michigan 48108, USA

**Keywords:** Adaptive radiation, section *Anisophyllum*, *Euphorbia*, Euphorbiaceae, Hawaiian Islands, subgenus *Chamaesyce*, taxon cycle

## Abstract

**Aim** The taxon cycle hypothesis describes the cyclic movement of taxa during range expansion and contraction, accompanied by an evolutionary shift from open and often coastal vegetation to closed, and often inland forest vegetation in island systems. The Hawaiian Archipelago is an ideal system to test this hypothesis given the linear fashion of island formation and a relatively well-understood geological history.

**Location** Hawaiian Islands.

**Methods** We sampled 153 individuals in 15 of the 16 native species of Hawaiian *Euphorbia* section *Anisophyllum* on six major Hawaiian Islands, plus 11 New World close relatives, to elucidate the biogeographic movement of this lineage along the Hawaiian island chain. We used a concatenated chloroplast DNA data set of more than eight kilobases in aligned length and applied maximum likelihood and Bayesian inference for phylogenetic reconstruction. Connectivity among islands and habitat types was estimated using BayesTraits. Age and phylogeographic patterns were co-estimated using BEAST. In addition, we used nuclear ribosomal ITS and the low-copy genes *LEAFY* and *G3pdhC* to investigate the reticulate relationships within this radiation.

**Results** We estimate that Hawaiian *Euphorbia* first arrived on Kauai or Niihau ca. 5 million years ago and subsequently diverged into 16 species on all major Hawaiian Islands. During this process *Euphorbia* dispersed from older to younger islands in a stepping-stone fashion through open, dispersal-prone habitats. Taxa that occupy closed vegetation on Kauai and Oahu evolved *in situ* from open vegetation taxa of the same island. Consequently, widespread species tend to occupy habitats with open vegetation, whereas single island endemic species predominantly occur in habitats with closed canopy and are only found on the two oldest islands of Kauai and Oahu.

**Main conclusions** The spatial and temporal patterns of dispersal and range shifts in Hawaiian *Euphorbia* support an intra-volcanic-archipelago version of the taxon cycle hypothesis.

## Introduction

The taxon cycle hypothesis describes the cyclic pattern of range expansion, contraction, and habitat shifts that occur in island archipelagos (Wilson, 1959, Wilson 1961; Ricklefs & Cox, 1978; Ricklefs & Bermingham, 2002). According to this hypothesis, taxa that colonize oceanic islands have inherently high dispersal abilities and are typically adapted to disturbed habitats. After the initial colonization, some populations expand to habitats that have closed forest canopies and are less susceptible to disturbance. During this transition, selection favors survival and competitive ability under closed vegetation rather than the dispersal ability that facilitated the initial colonization. Species that remain in the lowland, drier and more disturbed habitats retain their dispersal ability and continue to colonize new islands, whereas species moved to forests with closed vegetation are increasingly specialized and isolated, with a tendency to become narrow endemics.

The taxon cycle hypothesis was originally proposed to explain patterns observed among the ants of Melanesia (Wilson, 1959, Wilson 1961). It has been further developed in birds in the Lesser Antilles (Ricklefs & Cox, 1972; Ricklefs & Cox, 1978;Ricklefs & Bermingham, 1999), and supported by genetic evidence in birds of the Indo-Pacific (Jønsson *et al.*, 2014) and in ants across the Old World (Economo *et al*, 2015). Volcanic hot spot island archipelagos, such as the Hawaiian Islands, present an ideal system to test the taxon cycle hypothesis. The recent, relatively well-understood geological history makes the dynamics of taxon movement more tractable than in older, more complicated continental island systems.

There are 17 species of *Euphorbia* L. (Euphorbiaceae) native to the Hawaiian Islands. Sixteen of them form a clade within section *Anisophyllum*Roep. of *Euphorbia* subgenus *Chamaesyce* Raf. (hereafter referred to as Hawaiian *Euphorbia*; Yang & Berry, 2011). The other species, *E. haeleeleana* Herbst, represents a separate colonization event belonging to *Euphorbia* subgenus *Euphorbia* (Dorsey *et al.*, 2013) and is outside the scope of this study. The 16 Hawaiian *Euphorbia* species occur in all major island habitats, from coastal strand to dry forests, wet forests and bogs, and range in habit from subshrubs and shrubs to trees ten meters tall (Fig. 1). Four of these species have two or more recognized varieties. Ten species are endemic to a single major island, while the remainder are known from two or more major islands (Table 1). Previous phylogenetic studies suggest that Hawaiian *Euphorbia* originated through allopolyploidy, with their closest relatives being small herbs occurring in dry, warm, and exposed habitats from the New World, including *E. stictospora* Engelm., *E. velleriflora* (Klotzsch & Garcke) Boiss., *E. mendezii*Boiss., *E. leucantha* (Klotzsch & Garcke) Boiss., and *E. cinerascens* Engelm (Fig. 1f; Yang & Berry, 2011). Given the overlapping distribution of the putative sister species in North America, the allopolyploidy event likely happened before longdistance dispersal to Hawaii.

**Figure 1.**
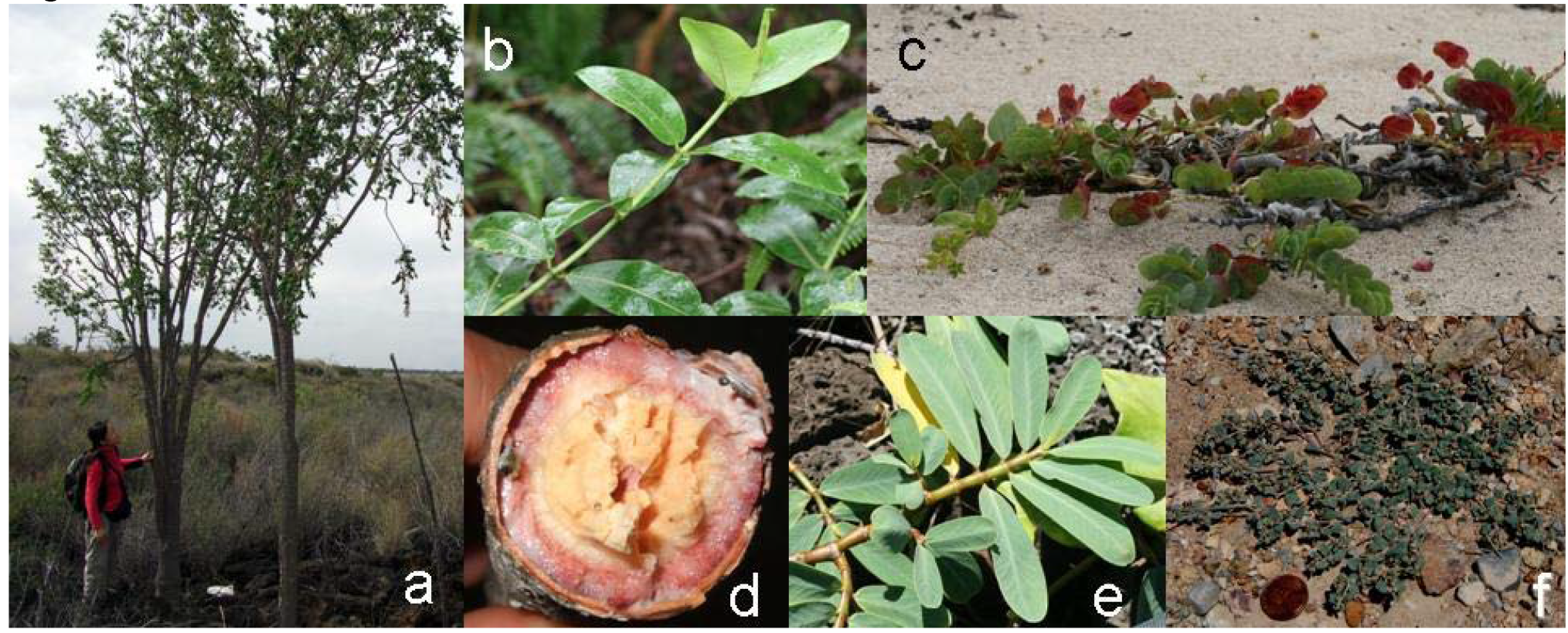
Hawaiian *Euphorbia* (a-e) and their closely related North American species (f). (a) *Euphorbia olowaluana*, a dry forest pioneer species on recently formed lava field, Hawaii; (b) *E. remyi* var. *remyi*, an ascending shrub in wet forest understory, Kauai; (c) *E. degeneri*, a prostrate subshrub on sandy beach, Oahu; (d) soft and fleshy woody stem of *E. celastroides* var. *kaenana;* (e) *E. celastroides* var. *kaenana*, a prostrate shrub, Oahu; (f) *E. cinerascens*, a small, prostrate perennial herb native to deserts in southern United States and northeastern Mexico (see coin in the lower left corner for scale).

In this study, we use multiple chloroplast and nuclear markers to reconstruct the history of radiation in Hawaiian *Euphorbia*. We adopt a modified classification of the taxon cycle stages (Wilson, 1959; Ricklefs & Cox, 1978), with stage I consisting of species that are undifferentiated and that occupy multiple islands (“widespread”), stage II consisting of widespread species with considerable intraspecific differentiation and recognized varieties, and stage III consisting of single island endemics. If Hawaiian *Euphorbia* fits the taxon cycle hypothesis, we predict that:

1. Hawaiian *Euphorbia* colonized younger islands successively from older islands.
2. Habitat connectivity is high among open vegetation and low among closed vegetation.
3. Stage I species predominantly occupy open vegetation and occur on all major islands; stage III species are most diverse in closed vegetation of older islands; whereas stage II species occur in a variety of habitat types on most islands.

## MATERIALS AND METHODS

### Taxon sampling

A total of 153 Hawaiian DNA accessions representing 15 of the 16 species of Hawaiian *Euphorbia* were included in this study. These samples covered six major Hawaiian Islands: Kauai, Oahu, Molokai, Maui, Lanai and Hawaii. The islands of Molokai, Maui, and Lanai together form the Maui Nui island group, given their past land connection as recent as the last interglacial period (Price & Elliott-Fisk, 2004). Of the 153 DNA accessions, 125 were obtained from the Hawaiian Plant DNA Library (Morden *et al.*, 1996; Randell & Morden, 1999), complemented by 18 additional samples newly collected from the field or cultivated sources (Appendix S1). Forty-three DNA accessions in the DNA Library were collected by M.J. Sporck-Koehler and L. Sack, accompanied by some of Hawaii's most experienced field botanists (see Acknowledgements) as part of an ecophysiological study of the Hawaiian *Euphorbia* (Sporck, 2011); permits for many of the species were limited to less than ten leaves per plant, with vouchers not permitted for State and Federally listed endangered or very rare and extremely vulnerable taxa. We described in Appendix S1 alternative voucher specimens representing the same population and/or additional locality information for source populations.

The resulting infraspecific sampling ranged from between one and 23 accessions per species. For species such as *E. deppeana*, which is found in only one wild population with ca. 50 individuals in total, only one accession was included; on the opposite side of the spectrum, *E. celastroides* var. *amplectens* and *E. degeneri* are both found on all major Hawaiian Islands, and 12 and 13 accessions were included, respectively, representing multiple populations from different islands. To distinguish different accessions of the same taxon, we included DNA numbers following taxon names for all the ingroup Hawaiian *Euphorbia* in the text. In addition, eleven closely related North American species were selected for outgroup comparison (Yang & Berry, 2011).

### Laboratory procedures

Genomic DNA extraction, plus PCR amplification and sequencing of both ITS and cpDNA followed the protocols in Yang & Berry (2011). A total of seven chloroplast (cpDNA) noncoding regions were sequenced: *rpl14-rpl36* spacer, *psbB-psbH* spacer, *atpI-atpH* spacer, *psbD-trnT*spacer, *trnH-psbA* spacer, *rpl16* intron, and the *trnL-F*region. For the ITS region, sequences with continuous superimposed peaks were excluded. Two of these excluded PCR products, *E. celastroides* var. *kaenana* 5840 and *E. kuwaleana* 5700 were cloned following the protocol of Yang & Berry (2011) to evaluate allelic variation. The second intron of the nuclear low-copy gene *LEAFY* and intron of *glyceraldehyde 3-phosphate dehydrogenase subunit C (G3pdhC)* were PCR amplified and cloned following the protocol in Yang & Berry (2011), except that at least 24 clones from each PCR product were sequenced. Copy-specific primer pairs were designed for both markers and at least eight clones were sequenced from each copy-specific PCR reaction. See Supplementary Methods in Appendix S2 for primers used, and PCR and cloning procedures using copy-specific primers.

### Phylogenetic analysis

Preliminary phylogenetic analyses using individual cpDNA non-coding regions detected three short inversions (Supplementary Methods, Appendix S2). All three inversions were reversed and complemented before concatenating all seven cpDNA regions into the first character set of the cpDNA matrix. Indels were scored following the simple gap-coding criterion (Simmons & Ochoterena, 2000) in SeqState v1.4.1 (Müller, 2006) and were treated as the second character set of the cpDNA matrix.

Bayesian inference was conducted in MrBayes v3.1.2 (Huelsenbeck & Ronquist, 2001; Ronquist & Huelsenbeck, 2003). Two independent runs of four chains each (three heated, one cold), starting from random trees, using a temperature of 0.2, were run for 10 million generations, using the model GTR+I+γ selected by AIC in MrModeltest v2.3 (Nylander, 2004). Trees were sampled every 1,000 generations. Parameters were unlinked between the two partitions except tree topologies. The binary indels were subject to “rates=gamma”. In addition, a branch length prior “brlenspr=unconstrained:exponential(100.0)” was applied to the nucleotide partition to prevent unrealistically long branches (Marshall, 2010). Diagnostic parameters were visually examined in the program Tracer v1.5 (Rambaut & Drummond, 2007) to verify stationary status. Trees sampled from the first 1 million generations were discarded as burn-in, and the remaining 18,002 trees were used to compute the majority rule consensus (MCC) tree.

Maximum likelihood (ML) analysis was carried out using RAxML v7.4.2 (Stamatakis, 2006), partitioning nucleotides vs. indels. The nucleotide substitution model was set to GTR+γ, 500 rapid bootstrap replicates were performed, followed by a thorough search for the best tree.

### Molecular dating

The Hawaiian island chain was formed by the Pacific plate moving northwestward over a fixed hot spot (Carson & Clague, 1995). We assumed that a new island was colonized soon after it emerged (Fleischer *et al*, 1998), and that given the extremely small colonizing population, deep divergence from ancestral polymorphisms in the colonizing population is highly unlikely. We cross-validated these two assumptions with a preliminary analysis estimating the stem ages of Maui Nui and Hawaiian clades by constraining the stem age of the oldest Oahu clade on the cpDNA data set with the time of full development of the Waianae Mountains (a normal prior with mean 3.86 my, standard deviation 0.089 my; Sherrod *et al.*, 2007; Lerner *et al.*, 2011).

A final analysis was carried out by applying the following age constraints to the cpDNA data set: (1) 3.86 ± 0.089 my for the stem age of the Oahu clade; and (2) 2.14 ± 0.117 my for the stem age of Maui Nui clades (the age of Penguin Bank, which formed the past land connection between Oahu and Maui Nui; Carson & Clague, 1995; Price & Elliott-Fisk, 2004; Sherrod *et al.*, 2007; Lerner *et al.*, 2011). The analysis was performed in BEAST v1.7.4 (Drummond *et al.*, 2012), using the concatenated cpDNA data set without coding indels. The substitution model HKY+I+**γ** was applied as selected by jModeltest v0.1.1 (Posada, 2008), with an uncorrelated lognormal relaxed clock and a pure-birth Yule model. Four independent runs of 60 million generations were carried out, sampling every 10,000 generations starting from a random starting tree. Convergence diagnose parameters were visualized in Tracer, and trees sampled from the first six million generations were discarded as burn-in. A MCC tree was calculated in TreeAnnotator v1.7.4.

### Phylogeographic reconstruction

Discrete phylogeographic analysis (Lemey *et al.*, 2009) was used to reconstruct the pattern of dispersal along the island chain from the cpDNA data set. Phylogeographic analysis was carried out in BEAST using two independent continuous-time Markov chains (CTMCs) by manually editing the xml file generated by BEAUti from the previous molecular dating analysis following Lemey *et al*. (2009). Most recent common ancestor (MRCA) of all Hawaiian accessions was set to Kauai according to molecular dating results. Convergence diagnostic parameters were visualized in Tracer, and the first six million generations were discarded as burn-in.

### Assignment of habitat types

We categorized coastal strand, scrub, and dry forest as “open vegetation”. These vegetation types are either fully exposed or have relatively open forest canopy coverage, and are generally relatively low in elevation, although the upper elevation limit of lowland dry forest varies from 150 to 1,500 meters depending on the island and the direction of the slope, and *E. olowaluana* occurs in montane dry forests from 700 to 2,800 m in elevation on Hawaii (Gagné & Cuddihy, 1990; Koutnik & Huft, 1990). Both mesic and wet forests have a closed forest canopy and were categorized as “closed vegetation”. Both generally occur at relatively high elevations. Montane bogs, although not protected by a closed forest canopy, are specialized forest openings that are surrounded by wet or mesic forests, and were categorized as closed vegetation.

### Estimating transition frequencies and connectivity among islands and habitat types

Each individual plant was assigned one of the six ranges according to their collection locality: Kauai open, Kauai closed, Oahu open, Oahu closed, Maui Nui open, or Hawaii open (no *Euphorbia* is native to closed vegetation on Maui Nui or Hawaii). For taxa that occur in more than one of these six ranges, we assigned a range for each individual according to its collection locality. The rates of transition among these six ranges were estimated using BayesTraits v1.0 (Pagel & Meade, 2007) by sampling from 1,000 random post burn-in trees from all four runs in the BEAST dating analysis, with outgroups coded as range unknown. Preliminary short reverse-jump Markov chain Monte Carlo (rjMCMC) runs were carried out to evaluate acceptance rates. Three independent final rjMCMC runs were carried out each with 50 million generations, sampling every 1,000 generations, with rate deviation set to 0.5 to optimize the acceptance rate, and a uniform hyperprior between 0 and 5 to seed the exponential prior. Results were visualized in Tracer, and rates sampled from the first 10 million generations were discarded as burn-in.

## RESULTS

### Extensive discordance between cpDNA and ITS in Hawaiian *Euphorbia*

We obtained sequences of all seven chloroplast non-coding regions from each of the 164 DNA accessions included in this study (Table S2.1 and Appendix S3). The aligned matrix was 8,278 bp in length. Branch lengths within Hawaiian *Euphorbia* were much shorter compared to the outgroup species (Fig. 2, upper left corner). Of the 13 species for which multiple individuals were represented in our sampling, 11 are either para-or polyphyletic according to the cpDNA tree, with the only exceptions being *E. herbstii* and *E. kuwaleana*, two rare species endemic to Oahu (Fig. 2). Despite being highly non-monophyletic at the species level, the phylogeny displayed strong geographical structuring. Monophyly of Hawaiian *Euphorbia* was well supported (PP=1 and BS=100; Fig. 2 & S2.2). A Kauai clade was sister to the rest of the Hawaiian *Euphorbia*, within which there are three well-supported Oahu clades (PP=1 and BS=78, 97 and 100 respectively). Among the three Oahu clades, the largest one (Oahu clade 1) had three well-supported Maui Nui clades (PP=1 and BS=62, 96, and 97 respectively) and one well-supported Hawaii clade (PP=1 and BS=97) nested in it. The only Kauai members in the Oahu clade were a small clade of *E. degeneri* nested in Maui Nui clade 2, which is a coastal strand species that occurs on all main islands.

**Figure 2.**
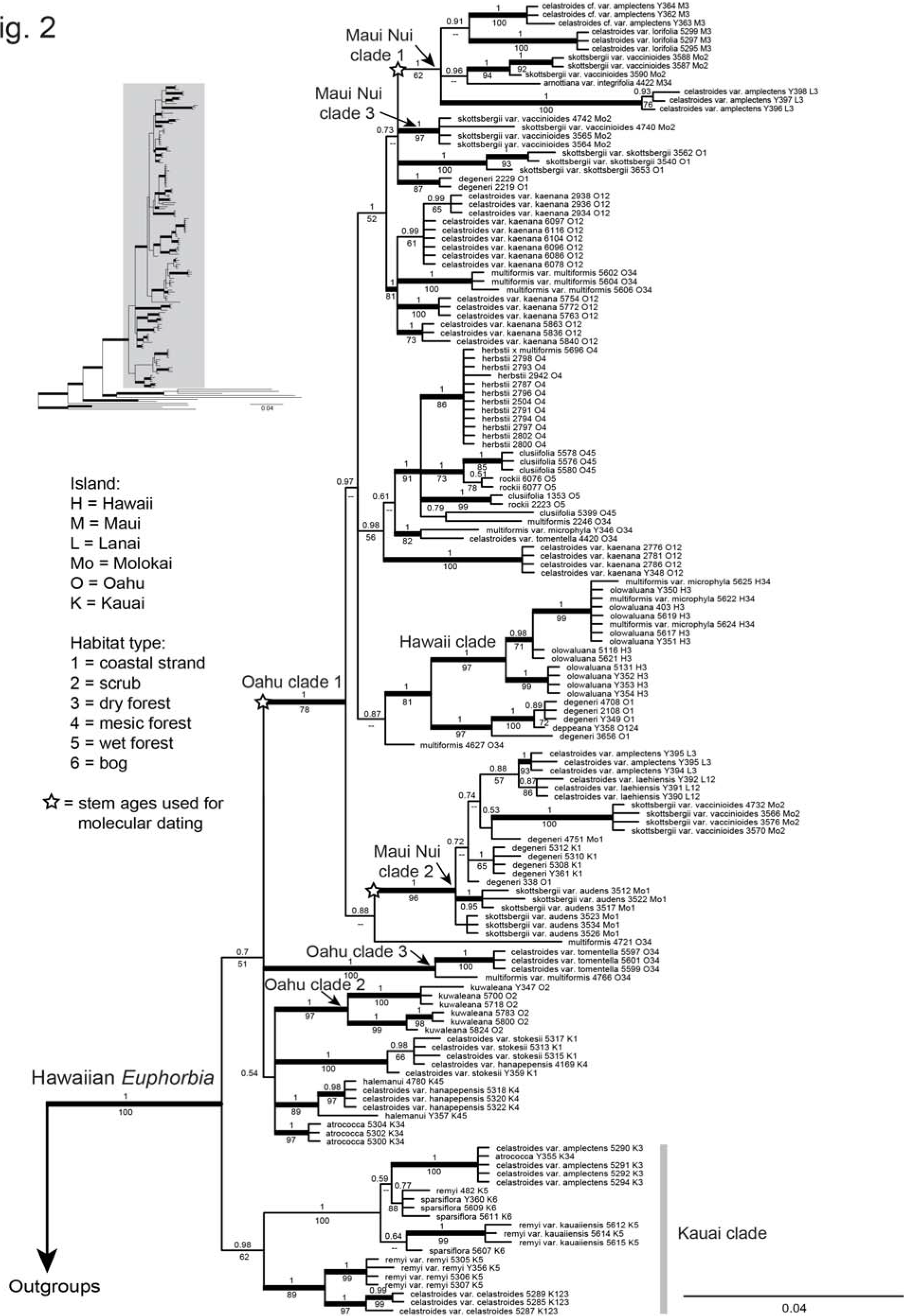
Majority rule consensus tree recovered from Bayesian analysis of cpDNA data. Numbers above the branches are Bayesian posterior probabilities (PP) and numbers below the branches are maximum parsimony bootstrap percentages (BS). Branch length scale is on lower right. Thick branches represent strongly supported clades with PP ≥ 0.95 and BS ≥ 70. Outgroups were removed on the main graph, with the full tree in the upper left corner. Following each taxon name is the DNA accession number, island initials for the individual, and vegetation type for the taxon.

Similar to the cpDNA phylogeny, the nuclear ITS tree highly supported the monophyly of Hawaiian *Euphorbia* (PP=1 and BS=100; Fig. S2.2a), as well as a Kauai grade with Oahu and younger island accessions nested in it. All ingroup ITS sequences have ten or more nucleotide positions showing superimposed peaks, which is much higher compared to outgroups. In addition, 18 of the ingroup accessions showed continuously superimposed peaks, likely from allele length variation and were excluded from the alignment. Cloning of *E. kuwaleana* 5700 and *E. celastroides* var. *kaenana* 5840 revealed many divergent alleles (Fig. S2.2a). Although evolution of the ITS region was highly dynamic, there are nonetheless a number of well-supported clades. Most of these clades occupied similar habitat types on a single island or Oahu+Maui Nui.

Both the low-copy nuclear genes *LEAFY* and *G3pdhC* showed increased copy numbers among Hawaiian taxa compared to outgroups, but the resolution within each copy was low (Fig. S2.2b). Four copies of *LEAFY* were recovered, of them only one copy was detected from the known outgroups. Similarly, six copies of *G3pdhC* were detected in Hawaiian *Euphorbia*, among which three had a clear association with known outgroup taxa.

### Molecular dating suggested a Kauai/Niihau origin of Hawaiian *Euphorbia*

We used cpDNA only for dating and phylogeographic analyses to track dispersal via seed dispersal or vegetative fragments. Using island age for molecular dating can potentially be biased by delayed arrival long after island formation, multiple dispersal events, local extinction, and ancestral polymorphism. Another consideration is that at the time Kauai formed ca. 5 mya, the adjacent island of Niihau was of similar size and prominence (Price & Clague, 2002). To cross-validate our assumptions and their potential caveats, a preliminary analysis was carried out constraining the stem age of Oahu clade 1, the most diverse and well supported Oahu clade, by the date at which the Waianae Mountains of Oahu formed (Fig. 2; 3.86 ± 0.089 my; Sherrod *et al.*, 2007; Lerner *et al.*, 2011). The resulting estimate for the median stem age of Maui Nui clade 1 was 2.5 my (95% credibility interval 1.6-3.3 my), Maui Nui clade 2 was 2.4 (1.5–3.2) my, and that for Maui Nui clade 3 was 1.4 (0.7-2.1) my. Both Maui Nui clades 1 and 2 had diversified on Maui Nui, and both had stem ages very similar to the age of Maui Nui (ca. 2.1 my; Sherrod et al., 2007; Lerner et al., 2011), whereas Maui Nui clade 3 is a much smaller and younger clade including only a single coastal taxon, and probably represented a more recent dispersal event. As for the Hawaii clade, both its stem age (1.9 my; 1. 1–2.9 my) and crown age (1.3 my; 0.7-2.1 my) were much older than the age of the island of Hawaii (≈0.59 my; Sherrod *et al*, 2007; Lerner *et al*, 2011). Both taxa in the Hawaii clade, *E. multiformis* var. *microphylla* and *E. olowaluana*, also occur on Maui Nui (Koutnik, 1987). It is likely that the “Hawaii clade” diverged on Maui Nui before dispersing to Hawaii.

Based on our cross-validation of dating points, our final molecular dating analysis constrained the stem age of Oahu clade 1 with the age of Oahu (3.86 ± 0.089 my) and Maui Nui clade 1 and 2 with the age of Maui Nui (2.14 ± 0.117 my). The resulting stem age of Hawaiian *Euphorbia* was estimated at 5.0 (4.1-6.3) my, around the time that Kauai and Niihau formed (ca. 5.1 mya; Fig. S2.3).

### Phylogeographic reconstruction supports successive island colonization

By co-estimating geographic distribution and tree topology, the resulting MCC tree from the phylogeographic reconstruction (Fig. 3) weakly supported Oahu clades 1, 2, and 3 as monophyletic (PP=0.46), as well as Maui Nui clades 1 and 3 as monophyletic (PP=0.49), instead of being non-monophyletic in the MCC tree of molecular dating alone (Fig. S2.3). All analyses from cpDNA using RAxML, MrBayes, as well as molecular dating and phylogeographic reconstruction strongly support a general trend of successive island colonization from older to younger islands, despite the disagreements in weakly supported nodes.

**Figure 3.**
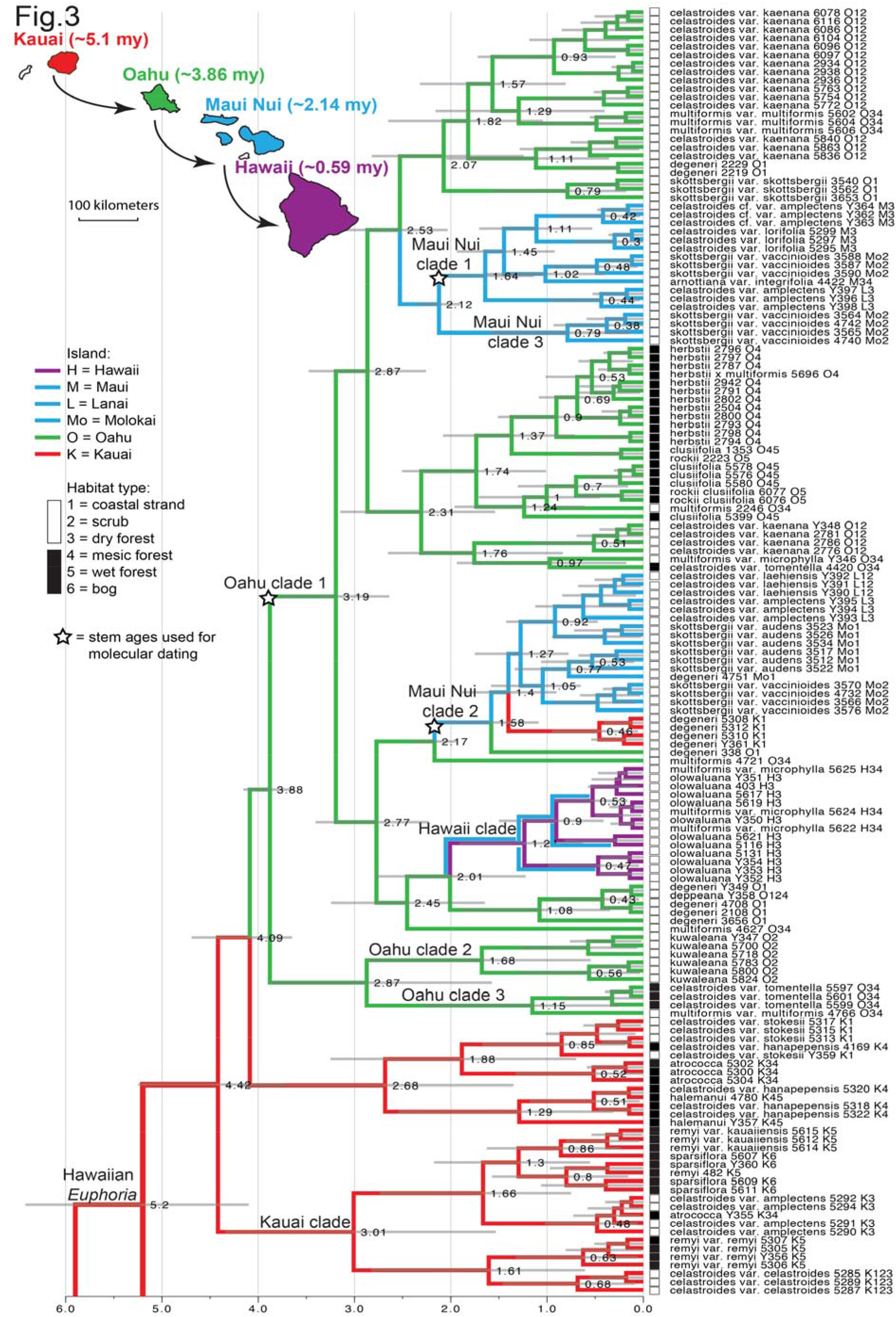
Maximum clade credibility (MCC) tree recovered from BEAST phylogeographic analysis. Node labels are mean ages, and node bars are 95% highest posterior density (HPD) intervals. Outgroups are not shown. Following each taxon name is the DNA accession number, island initials for the individual, and vegetation type for the taxon. Superimposed blue lines on the Hawaii clade indicates the most likely scenario inferred from the clade age and historical distribution of *E. olowaluana* on Maui. Map in the upper left corner shows the inferred dominant pattern of dispersal among islands.

### BayesTraits analysis supports contrasting connectivity between open and closed vegetation

BayesTraits analysis was consistent with high habitat connectivity within islands and among open habitats across islands, but low connectivity among closed habitats (Table 2, Fig. 4). The most frequent transitions occurred between open and closed Kauai, the oldest of the four islands; transitions were also frequent among open Kauai, open Oahu and open Maui Nui. In contrast to the frequent exchange among open habitats, there was little exchange between closed Kauai and closed Oahu (Table 2, Fig. 4). Among all the six ranges, open Hawaii was most isolated and involved the fewest transitions, consistent with its recent origin. Open Oahu is the most connected, serving as stepping stone between Kauai and the two younger island groups of Maui Nui and Hawaii. The ancestral range was reconstructed as 19.3% open Kauai, 32.5% closed Kauai, 17.1% open Oahu, 11.7% closed Oahu, 11.3% open Maui, and 8.1% open Hawaii.

**Figure 4.**
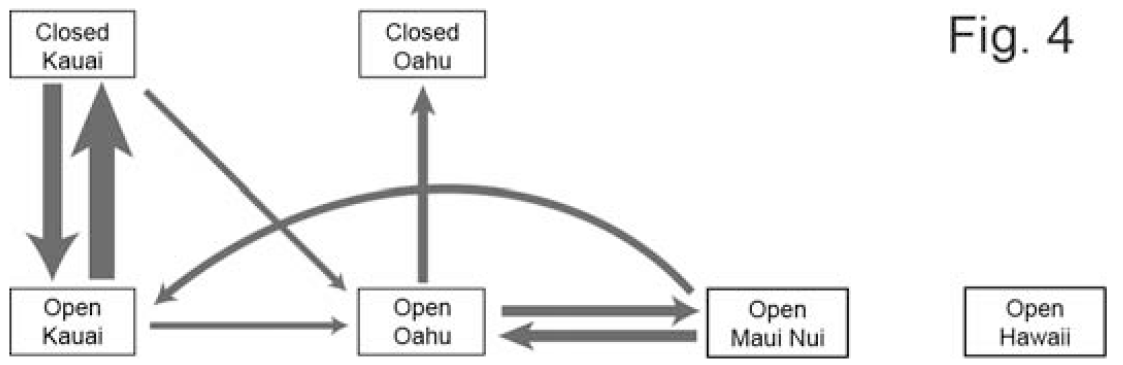
Transition rates among all six ranges where Hawaiian *Euphorbia* occurs. Thickness of arrow is proportional to the median value of the transition rate. For clarity, rates lower than 0.03 are now shown.

### Distribution of taxon richness across islands and habitats

The number of overall species per island was highest in Oahu, and decreased towards both older and younger islands, showing a humped trend (Fig. 5a). Stage I species were most numerous on Maui Nui; stage II species were approximately equally numerous on all four islands; and stage III species were only found on Kauai and Oahu, the two oldest islands, and absent from the two younger island groups. Alternatively, breaking down infraspecific taxa by endemic to one island vs. on two or more islands (Fig. 5c) showed a similar trend, with the number of endemic taxa highest on Kauai, and the number of widespread taxa peaking on Maui Nui.

**Figure 5.**
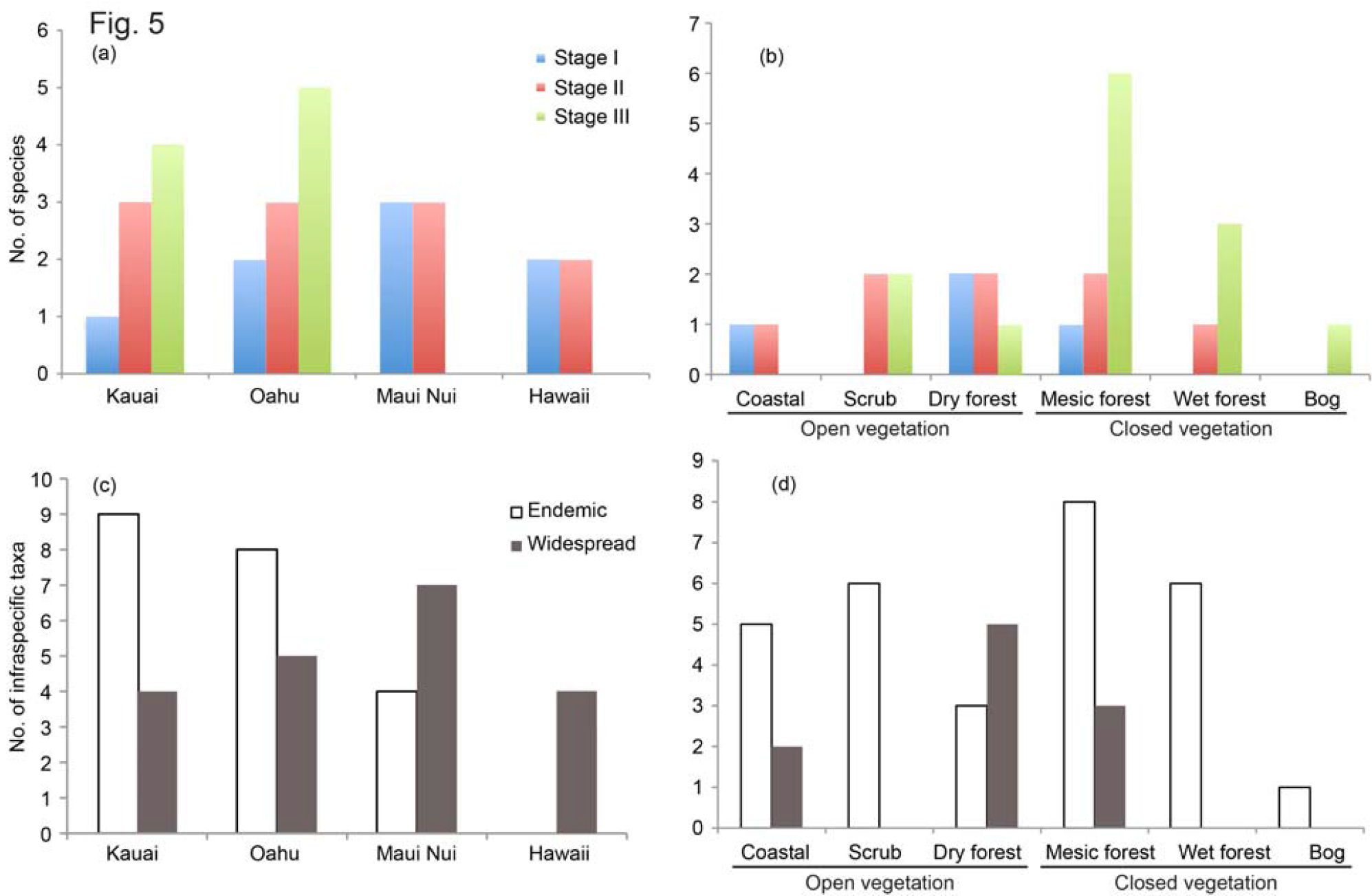
Number of species in each taxon cycle stage (a) on each major island, and (b) in each habitat type. Number of taxon including varieties that are endemic to a single major island or more widespread (c) on each major island, and (d) in each habitat type.

The species-habitat plot (Fig. 5b) showed that stage I species occurred mostly in open habitats (coastal strand and dry forest), stage II species occurred in both open and closed habitats, and stage III species mostly occurred in closed habitats. Similarly, when plotted by infraspecific taxa, widespread taxa were most numerous in dry forest, whereas endemic taxa were most numerous in mesic forest.

## DISCUSSION

### Rapid, reticulate radiation of Hawaiian *Euphorbia* from an allopolyploid ancestor

Our analyses supported radiation from a single colonization through allopolyploidy. Both of the nuclear low-copy genes examined in this study, *LEAFY* and *G3pdhC* had increased copy numbers in Hawaiian *Euphorbia* compared to outgroups. Two of the four copies detected in *LEAFY*, and three of the six copies detected in *G3pdhC*were not associated with known outgroups previously identified using ITS and chloroplast markers, consistent with previous results from cloning another nuclear low-copy gene, *EMB2765* (Yang & Berry, 2011). Additional support for the allopolyploid origin of Hawaiian *Euphorbia* comes from chromosome numbers: while the outgroup species *E. cinerascens* and *E. stictospora* have chromosome counts of 2n=12 and 2n=32 respectively (Urbatsch *et al.*, 1975), all four Hawaiian species surveyed so far have 2n=38 (E. *arnottiana, E. celastroides, E. clusiifolia*, and *E. multiformis*, Carr, 1985).

Following the putative initial allopolyploidy event, Hawaiian *Euphorbia* diversified rapidly with reticulation. All species that occur in open habitats are highly polyphyletic according to cpDNA, with the only exception being *E. kuwaleana*, a rare species that occurs on Oahu. Of the species occupying open vegetation, *E. degeneri* has distinctive sessile leaves that are rounded in shape and fold upwards and is specialized in coastal beach habitats (Fig. 1c; Koutnik, 1987). We included multiple Kauai, Oahu, and Maui Nui accessions of *E. degeneri*, and they are placed by cpDNA in multiple, separate clades within Oahu clade 1, and by ITS in a polytomy consisting mostly of open Oahu and younger island accessions. Given its distinctiveness in both morphology and habitat, its polyphyly in the cpDNA tree, and the lack of resolution in the ITS tree, what we recognize morphologically and ecologically as *E. degeneri* is likely a result of reticulate evolution and/or rapid diversification lacking species monophyly.

A second highly polyphyletic species, *E. celastroides* (Fig. 1d–e), is more variable in both morphology and habitats and has eight recognized varieties (Table 1). Varieties of *E. celastroides* can be prostrate or upright, with leaf surfaces varying from glabrous to papillate, and the cyathia range from solitary to multiple. Each variety occupies one or more habitat types from coastal strand to mesic forest, and may be either endemic to a single island or more widespread. Of the varieties, *E. celastroides* var. *kaenana* is endemic to the northwestern corner of Oahu. Despite its restricted distribution, it is polyphyletic and shows intermixture with *E. multiformis* from the same island in the cpDNA phylogeny (Fig. 2), whereas ITS places all accessions of *E. celastroides* var. *kaenana* in a polytomy with stage I and other stage II species *(E. degeneri, E. multiformis, E. skottsbergii*, and other varieties of *E. celastroides*, Fig. S2.2a). Despite being highly variable, *E. celastroides* is morphologically recognizable, with entire, distichous leaves that are oblong to obovate in shape (Fig. 1e). It can be distinguished from the vegetatively similar *E. multiformis*, also a widespread stage II species, by its erect fruits and appressed cyathial glands, as opposed to recurved fruits and protruding glands in *E. multiformis* (Koutnick 1985).

In addition to presumed older reticulations, we also found evidence for more recent hybridization events. *Euphorbia multiformis* var. *microphylla* 5622 and 5624 were collected from the Pohakuloa Training Area of Hawaii, and they share an almost identical cpDNA haplotype with *E. olowaluana* accessions that came from the same area (Fig. 2). In the ITS phylogeny, however, both *E. multiformis* var. *microphylla* 5622 and 5624 are not part of a monophyletic *E. olowaluana* (Fig. S2.2a).

Similar patterns of taxon non-monophyly are also found in other endemic Hawaiian plant lineages when multiple accessions are sampled and show evidence of reticulate evolution and rapid diversification. These include *Scaevola* (Goodeniaceae; Howarth & Baum, 2005), *Plantago* (Plantaginaceae; Dunbar-Co *et al.*, 2008), *Metrosideros* (Myrtaceae; Percy *et al.*, 2008), *Pittosporum* (Pittosporaceae; Bacon *et al*, 2011), and *Bidens* (Asteraceae; Knope *et al*, 2012). Together these examples caution against using single representative samples per species, or relying on cpDNA and/or ITS as the sole source for studying recent, rapid radiations.

### Kauai origin and stepping-stone dispersal from older to younger islands

Overall, our analyses suggest that Hawaiian *Euphorbia* first colonized Kauai or Niihau, then Oahu, Maui Nui, and finally Hawaii, following the “progression rule” from older to younger islands (Hennig, 1966; Funk & Wagner, 1995), with at least one dispersal event in the reverse direction through a stage I species.

Our BayesTraits analysis with outgroup ranges coded as “unknown” reconstructed the most likely origin of Hawaiian *Euphorbia* as Kauai (52%), followed by Oahu (29%; Table 2). Molecular dating using ages of Oahu and Maui Nui further supported a Kauai or Niihau origin of Hawaiian *Euphorbia*, given these two islands were of similar sizes 5 mya (Price & Clague, 2002). Our estimation of the stem age of Hawaiian *Euphorbia* at ca. 5 my is also consistent with previous molecular dating based on fossil evidence, which estimated the split between *E. hirta* and *E. humifusa* to be ca. 9 mya (Horn *et al.*, 2014), a split much deeper than the stem of Hawaiian *Euphorbia* (Yang & Berry, 2011).

Following the initial establishment in Kauai, dispersal from Kauai to Oahu occurred at least once (Fig. 3), with coastal Oahu serving as a hub and stepping stone for diversification in Oahu and to younger islands. There were at least two dispersal events from Oahu to Maui Nui, followed by back-dispersals from Maui Nui clade 2 to Kauai and probably also to Oahu, both involving the widespread coastal strand species *E. degeneri*. Although all individuals from Hawaii form a monophyletic clade, given that its crown age (0.63-1.91 mya) is significantly older than the age of the island (ca. 0.59 mya) and that both *E. multiformis* var. *microphylla* and *E. olowaluana* also occur on Maui Nui, the Hawaii clade likely split on Maui Nui before dispersing to Hawaii, as the superimposed blue lines on Fig. 3 indicate.

### High connectivity among open vegetation with *in situ* origin of clades of closed vegetation

Given that all outgroup species are from dry and disturbed habitats, the initial colonization of ancestral Hawaiian *Euphorbia* likely occurred on coastal Kauai. Although BayesTraits reconstructed the most likely root range as closed Kauai (32.8%), followed by coastal Kauai (19.3%, Table 2), reconstruction of the root as closed Kauai is probably an artifact of the low phylogenetic resolution at the base of Hawaiian *Euphorbia* and the movements towards closed Kauai during the relatively long period of occupation. This artifact likely also resulted in the reconstruction of an approximately equal transition rate from closed vs. open Kauai to open Oahu (Table 2 and Fig. 4). Given that all species in open vegetation on Kauai are widespread and all species in closed vegetation on Kauai are single-island endemics, the dispersal from Kauai to Oahu likely occurred through open habitats.

According to our BayesTraits reconstructions, open Oahu was the hub of the dispersal chain (Table 2 and Fig. 4), serving as the stepping-stone between open Kauai, closed Oahu and open Maui Nui. Very little connectivity was recovered to and from Hawaii, consistent with its recent geological history. On the other hand, transitions between closed Kauai and closed Oahu were close to zero, despite both ranges existing for more than three million years. The most frequent transition to closed Kauai was from open Kauai, and the most frequent transition to closed Oahu was from open Oahu. A similar pattern of “upslope migration” is also evident in Hawaiian *Artemisia* (Hobbs & Baldwin, 2013) and in flightless alpine moths in Hawaii and Maui (Medeiros & Gillespie, 2011). In Hawaiian violets, however, an nrITS phylogeny recovered a dry clade and a wet clade, each having species from multiple islands (Havran *et al.*, 2009). Since that study relied solely on the nrITS marker in a group of polyploid species, its results may have been biased (Marcussen *et al.*, 2012).

### The intra-volcanic-archipelago taxon cycles in Hawaiian *Euphorbia*

In addition to our findings of repeated dispersal across the island chain and movements toward closed habitats on individual islands, our findings of species and taxon richness across islands and habitat types also support the taxon cycle hypothesis. Widespread species with no recognized varieties (stage I) are distributed on all islands and only occur in open habitats (Fig. 5a–b). In contrast, all single island endemic species (stage III) occurred on Kauai and Oahu, the two oldest islands, and most of them occur in closed habitats. No stage III species occur on Maui or Hawaii, despite their larger sizes and higher elevations than the older islands. Therefore it appears that when a young island emerges, it is first colonized by stage I and stage II species. Stage III species only arise later through evolution of adaptation to forest understories.

Certain infraspecific taxa in Hawaiian *Euphorbia* are geographically and morphologically distinctive enough that it is sometimes unclear whether separate species should be recognized (see discussion in Koutnik 1985; Koutnik 1987; Koutnik & Huft, 1990). This kind of taxonomic uncertainty may reflect the process of increased isolation during the transition from stage II to stage III of a taxon cycle. We further broke down the distribution of infraspecific taxa by island and habit respectively (Fig. 5c–d). The trend is very similar to species vs. island and habitat relationships: single-island endemic taxa are most numerous on Kauai and Oahu, and absent from Hawaii, whereas the number of widespread taxa peaks on Maui Nui. Among habitat types, endemic taxa are most numerous in mesic forests, whereas widespread taxa are most numerous in dry forests.

Both dispersal ability and habitat specialization in Hawaiian *Euphorbia* appear to be associated with seed characters. Hawaiian *Euphorbia* most likely arrived from North America via tiny seeds that adhered to birds through a mucilaginous seed coat (Carlquist, 1966, Carlquist 1980a; Price & Wagner, 2004). A survey of mucilaginous seed coats across *Euphorbia* sect. *Anisophyllum* (Jordan & Hayden, 1992) showed that it is present in most mainland species as well as in *E. celastroides*, a stage II species and one of the most widespread members of Hawaiian *Euphorbia*. It is absent, however, in all four stage III Hawaiian species surveyed: *E. clusiifolia, E. halemanui, E. remyi*, and *E. rockii*. Interestingly, *E. degeneri*, a stage I species occurring on coastal strand of all major Hawaiian islands, lacks a mucilaginous seed coat. Instead, it is able to float in sea water (Carlquist, 1980b), which likely explains its coastal distribution and offers an alternative dispersal mechanism besides sticking to birds. In contrast, neither *E. celastroides* (stage II) nor *E. clusiifolia* (stage III) appear to have floating seeds (Carlquist, 1966). In addition to the difference in dispersal ability between species of different taxon cycle stages, stage III species such as *E. rockii* and *E. clusiifolia* have seeds about twice as large in diameter as stage I and stage II species. Such larger, non-sticky, non-buoyant seeds may enhance seedling survival in forest understory while simultaneously reducing their dispersal ability.

By being restricted to localized habitats and having much reduced dispersal ability, stage III species are likely more vulnerable to extinction, as predicted by the taxon cycle hypothesis. Of the six species and four additional varieties of Hawaiian *Euphorbia* that are federally listed as endangered, five species plus two additional varieties belong to stage III and predominantly occur in wet or mesic forest (*E. deppeana*, *E. eleanoriae*, *E. halemanui*, *E. herbstii*, *E. remyi* var. *kauaiensis, E. remyi* var. *remyi*, and *E. rockii)*, with *E. kuwaleana* being the only lowland scrub species listed, two remaining varieties are localized populations of stage II species *(E. celastroides* var. *kaenana* and *E. skottsbergii* var. *skottsbergii)* that occur in coastal habitats.

Although the cyclic pattern of dispersal and habitat shift seen in Hawaiian *Euphorbia* supports aspects of the taxon cycle hypothesis, there are also fundamental differences between the taxon cycles described by Wilson, 1959, Wilson 1961) and our intra-volcanic-archipelago version of the taxon cycle. First, Wilson described cyclic patterns at a larger scale, with repeated movement from the continent to an archipelago. In contrast, the pattern seen in Hawaiian *Euphorbia* occurred over smaller distances and times. Because of the highly isolated nature of the Hawaiian Archipelago, colonization from a continent is infrequent. Instead of “colonization waves” from the continent, colonization occurred in waves from older to younger islands, starting soon after a young island formed.

Secondly, in the original taxon cycle concept (Wilson, 1959, Wilson 1961), single island endemics formed from range contraction and local extinction from previously widespread species. In the Hawaiian Islands, however, ecological speciation occurred rapidly within islands such that populations occupying closed habitats speciated while the widespread species dispersed to the next new island, producing a high level of polyphyly among coastal taxa. Many stage II species and their infraspecific taxa are likely at various stages of moving to specialized habitats and becoming increasingly isolated. The parallel formation of new species in closed vegetation from lowland open vegetation on both Kauai and Oahu provides a promising case study for underlying mechanisms and processes driving ecological speciation.

Finally, the time series of island formation and erosion in volcanic island systems adds another dimension to the dynamics of taxon cycle (Whittaker *et al.*, 2008). This is evident from the hump-shaped curve of overall species versus island age relationship typical in volcanic island systems (Fig. 5), here showing a peak for Oahu. Stage I species lead the curve and peak on Maui Nui, stage II species had a more even distribution among major island groups, and stage III species occurred only on the two oldest islands. The total species number of Hawaiian lobelias, in contrast, peaked on Maui (Givnish et al., 2009), the second youngest major island. That pattern was likely due to an earlier arrival on a former tall island further up the island chain and/or more rapid divergence, making the Hawaiian lobelioids “saturated” with ca. 130 recognized species relative to the 16 of Hawaiian *Euphorbia*.

## Conclusions

Our analyses suggest that after initial colonization of Kauai or Niihau, Hawaiian *Euphorbia* moved in a stepping-stone fashion from older to younger islands through dry and disturbed open habitats, whereas species occupy closed vegetation evolved *in situ* on the older islands of Kauai and Oahu. During this process, widespread species become fragmented and gave rise to single island endemics.

The intra-volcanic-archipelago taxon cycles we found in Hawaiian *Euphorbia* merit testing in other lineages and island systems, as it has broad implications for lineage diversification in island systems in general. It predicts where tradeoffs occur among functional traits that are associated with dispersal versus establishment, the occurrence of species with high extinction risk, and offers testable hypothesis for ecological speciation.

## ACKNOWLEDGEMENTS

We thank the following people for help with field work: Christian Torres-Santana, Kenneth Wood, Larry Abbott, Ane Bakutis, Lala Bialic-Murphy, Pat Bily, Joanne Birch, Matt Burt, Molly Cavaleri, Marian Chau, Susan Ching, Margaret Clark, Vince Costello, Michelle Elmore, Steve Evens, Erin Foley, Julia Gustine, Kapua Kawelo, Matt Keir, Tobias Koehler, Joel Lau, Matthew Lurie, Kristen Nalani Mailheau, Steve Perlman, Allen Rietow, Dan Sailor, Wayne Souza, Natalia Tangalin and Mashuri Waite. The field work was facilitated by the following agencies: Hawaii Department of Land and Natural Resources, Division of Forestry and Wildlife, Oahu Army Natural Resources Program, the Center for Environmental Management of Military Lands, Colorado State University, Pohakuloa Training Area, Nature Conservancy of Hawaii, National Tropical Botanical Garden, and The Plant Extinction Prevention Program. We would like to thank Hank Oppenheimer for providing plant materials; Lauren Raz and Timothy Motley for allowing us to used their unpublished DNA samples; Evan Economo, Warren Wagner, and Jess Peirson for insightful discussions and help revising the manuscript. Funding was provided by the National Science Foundation through a Planetary Biodiversity Inventory award (DEB-0616533) to PEB and the National Tropical Botanical Garden through a McBryde Fellowship to PEB and YY.

## DATA ACCESSIBILITY

All DNA sequence data were deposited in GenBank (Appendix S1, to be deposited). All alignment files in NEXUS format are in Appendix S3.

## BIOSKETCH

Ya Yang is a postdoctoral associate working on plant systematics and functional phylogenomics (www.yangya.org). C.W.M. is a professor studying evolution and biogeography of the Hawaiian flora. M.J.S.-K. administers the State of Hawaii Rare Plant Program and studies plant ecophysiology and conservation.

**Table 1.** Distribution of the 16 Hawaiian *Euphorbia* species on six major Hawaiian Islands. Habitat types are sorted from wetter habitats generally at higher elevations to lower elevation and dryer ones, whereas ages of islands are ordered left to right from older to younger (Koutnik, 1987; Koutnik & Huft, 1990; Lorence & Wagner, 1996; Morden & Gregoritza, 2005). Taxa with an “*” are federally listed as endangered.

| Species | Varieties | Habit | Habitat | Taxon cycle stage | Kauai | Oahu | Maui Nui |  |  | Hawaii |
| --- | --- | --- | --- | --- | --- | --- | --- | --- | --- | --- |
|  |  |  |  |  |  |  | Molokai | Lanai | Maui |  |
| *sparsiflora* |  | Subshrub | Bog | III | X |  |  |  |  |  |
| *remyi* | *hanaleiensis* | Shrub | Wet forest | II | X |  |  |  |  |  |
| *remyi* | *kauaiensis* * | Shrub | Wet forest | II | X |  |  |  |  |  |
| *remyi* | *remyi* * | Shrub | Wet forest | II | X |  |  |  |  |  |
| *rockii* * |  | Shrub to small tree | Wet forest | III |  | X |  |  |  |  |
| *clusiifolia* |  | Shrub | Mesic to wet | III |  | X |  |  |  |  |
| *halemanui* * |  | Shrub | Mesic to wet | III | X |  |  |  |  |  |
| *celastroides* | *hanapehensis* | Shrub | Mesic forest | II | X |  |  |  |  |  |
| *eleanorae* * |  | Shrub | Mesic forest | III | X |  |  |  |  |  |
| *atrococca* |  | Shrub to small tree | Mesic forest | III | X |  |  |  |  |  |
| *herbstii* * |  | Tree | Mesic forest | III |  | X |  |  |  |  |
| *deppeana* * |  | Subshrub | Scrub to mesic forest | III |  | X |  |  |  |  |
| *celastroides* | *tomentella* | Shrub | Dry to mesic | II |  | X |  |  |  |  |
| *arnottiana* |  | Shrub | Dry to mesic | I |  | X |  |  | X |  |
| *multiformis* | *multiformis* | Shrub | Dry to mesic forests | II |  | X |  |  | X |  |
| *multiformis* | *microphylla* | Shrub | Dry to mesic forests | II | X | X | X |  | X | X |
| *olowaluana* |  | Tree | Dry forest and open sub-alpine forest | I |  |  |  |  | X | X |
| *celastroides* | *amplectens* | Shrub | Dry forest | II | X | X | X | X | X | X |
| *celastroides* | *lorifolia* | Shrub to small tree | Dry forest | II |  |  |  | X | X |  |
| *skottsbergii* | *vaccinioides* | Shrub | Scrub | II |  |  | X |  | X |  |
| *kuwaleana* * |  | Shrub | Scrub | III |  | X |  |  |  |  |
| *celastroides* | *celastroides* | Shrub | Coastal strand to dry forest | II | X |  |  |  |  |  |
| *celastroides* | *kaenana* * | Shrub | Coastal strand to scrub | II |  | X |  |  |  |  |
| *celastroides* | *laehiensis* | Shrub | Coastal strand to scrub | II |  |  |  | X | X |  |
| *celastroides* | *stokesii* | Shrub | Coastal strand | II | X |  | X |  |  |  |
| *degeneri* |  | Subshrub | Coastal strand | I | X | X | X |  | X | X |
| *skottsbergii* | *audens* | Shrub | Coastal strand | II |  |  | X |  |  |  |
| *skottsbergii* | *skottsbergii* * | Shrub | Coastal strand | II |  | X |  |  |  |  |

**Table 1.**
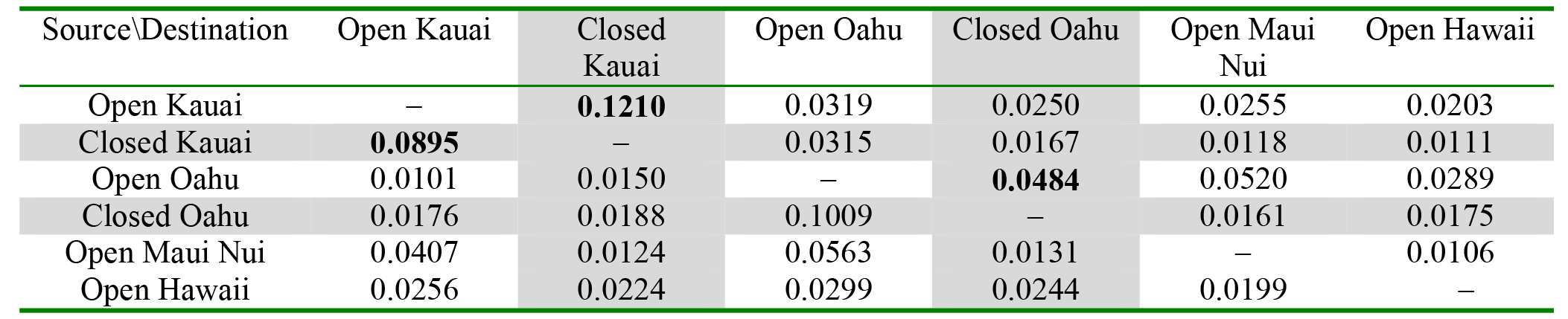
Median transition rates between ranges inferred from BayesTraits. Rates that are nonzero in >95% of the post-burn-in values are bold. Shaded cells involve either the source or destination range as being closed vegetation.

## SUPPORTING INFORMATION

Additional Supporting Information may be found in the online version of this article:

**Appendix S1** Collection, voucher and GenBank accession information.
**Appendix S2** Supplementary information
**Appendix S3** Alignments in NEXUS format and trees in Newick format.

